# Cost-efficient whole genome-sequencing using novel mostly natural sequencing-by-synthesis chemistry and open fluidics platform

**DOI:** 10.1101/2022.05.29.493900

**Authors:** Gilad Almogy, Mark Pratt, Florian Oberstrass, Linda Lee, Dan Mazur, Nate Beckett, Omer Barad, Ilya Soifer, Eddie Perelman, Yoav Etzioni, Martin Sosa, April Jung, Tyson Clark, Eliane Trepagnier, Gila Lithwick-Yanai, Sarah Pollock, Gil Hornung, Maya Levy, Matthew Coole, Tom Howd, Megan Shand, Yossi Farjoun, James Emery, Giles Hall, Samuel Lee, Takuto Sato, Ricky Magner, Sophie Low, Andrew Bernier, Bharathi Gandi, Jack Stohlman, Corey Nolet, Siobhan Donovan, Brendan Blumenstiel, Michelle Cipicchio, Sheila Dodge, Eric Banks, Niall Lennon, Stacey Gabriel, Doron Lipson

## Abstract

We introduce a massively parallel novel sequencing platform that combines an open flow cell design on a circular wafer with a large surface area and mostly natural nucleotides that allow optical end-point detection without reversible terminators. This platform enables sequencing billions of reads with longer read length (∼300bp) and fast runs times (<20hrs) with high base accuracy (Q30 > 85%), at a low cost of $1/Gb. We establish system performance by whole-genome sequencing of the Genome-In-A-Bottle reference samples HG001-7, demonstrating high accuracy for SNPs (99.6%) and Indels in homopolymers up to length 10 (96.4%) across the vast majority (>98%) of the defined high-confidence regions of these samples. We demonstrate scalability of the whole-genome sequencing workflow by sequencing an additional 224 selected samples from the 1000 Genomes project achieving high concordance with reference data.

## Introduction

Continuous advances in DNA sequencing over the last two decades have enabled cutting edge life sciences research by providing access to increasing amounts of genomic, transcriptomic, and epigenetic data across all fields of biology. Methods for massively parallel sequencing of clonally-amplified short DNA fragments, also known as second-generation sequencing, have reduced the sequencing cost of a whole human genome by over 6 orders of magnitude, from $3B estimated for the Human Genome Project^1^ to under $1000 per genome^2^ enabling a rapidly growing set of clinical applications ranging from carrier screening and prenatal testing to tumor profiling and early cancer detection^3–6^. However, sequencing cost reduction has stalled in recent years around the $6-10/Gb price point^2^. As a result, sequencing costs remain a critical bottleneck and limited funds often force tradeoffs between the breadth, depth, and frequency of genomic sequencing in the design of research and clinical assays. Resuming the push for sequencing cost reduction will drive the development of new genomic applications and the adoption of genomic diagnostics into the standard of care.

Here we introduce a new massively parallel sequencing-by-synthesis (SBS) approach which combines some of the most desirable traits of current short-read methodologies within a system providing significant scalability and dramatically lower cost. The first implementation of this approach produces approximately 10 billion reads per run, with a turnaround time of under 20hrs per run for 300bp reads, and with base quality similar to existing platforms (Q30 >85%), at a price of $1/Gb. We establish benchmark sequencing and germline variant calling performance on the standard Genome in a Bottle (GIAB)^7^ reference samples and demonstrate a scalable workflow by generating whole genome sequencing (WGS) data of over 200 previously characterized genomes.

### Sequencing Platform

To enable cost-effective high-scale DNA sequencing that is both superior to current methods and amenable to continuous improvement, we have designed a sequencing architecture that efficiently utilizes economical consumables and contains multiple degrees of freedom having significant headroom for additional scalability. Our system design features three main innovative components: a) *open fluidics and optics system*, b) *mostly natural sequencing chemistry*, and c) *neural network-enabled base-calling*. Combined, these innovations enable scalable, high-throughput DNA sequencing and significantly reduce the consumable cost of sequencing down to $1/Gb in the first implementation, with potential for even lower costs in the not distant future.

#### Open fluidics and optics system

The cost of sequencing is dominated by two consumables: the flow-cell and the sequencing reagents. Typical next generation sequencing (NGS) platforms utilize highly engineered flow-cells that both control reagent delivery and simultaneously function as part of the optical detection path. Selection of the fluidic channel dimensions within the flow-cell are a performance compromise between efficiency of reagent flow speed (deep channels) and reagent usage (shallow channels). Each surface of the flow-cell that is in the optical path must meet micron-scale mechanical tolerances consistent with high resolution microscopy. Furthermore, since a flow-cell is fed by common input lines, all of the fluidic channels shared between different reagents must be completely flushed between SBS steps to prevent carryover contamination, which is detrimental to data quality. In contrast, our system circumvents these design constraints by utilizing a circular 200mm silicon wafer as an “open flow-cell” with no consumable parts in the optical path. This wafer is patterned at micron scale generating a dense array of electrostatic landing pads to bind clonally amplified sequencing beads, which are separately produced by an automated emulsion PCR process (Fig. 1a). A spin-dispense system delivers reagents to the wafer by dispensing reagents from dedicated nozzles near the center of the rotating wafer and distributing the reagents rapidly and uniformly across the wafer by centrifugal force (Fig. 1b). This system has no shared fluidics between reagents that must be purged between cycles to avoid contamination. Because reagents are delivered inertially, a very uniform and thin layer (about 10 microns) is created on the surface, routinely achieving high active reagent utilization. Optical measurement of the entire surface is performed during rotation of the wafer in a continuous process, analogous to reading a compact disc (Fig. 1b).

**Figure 1:**
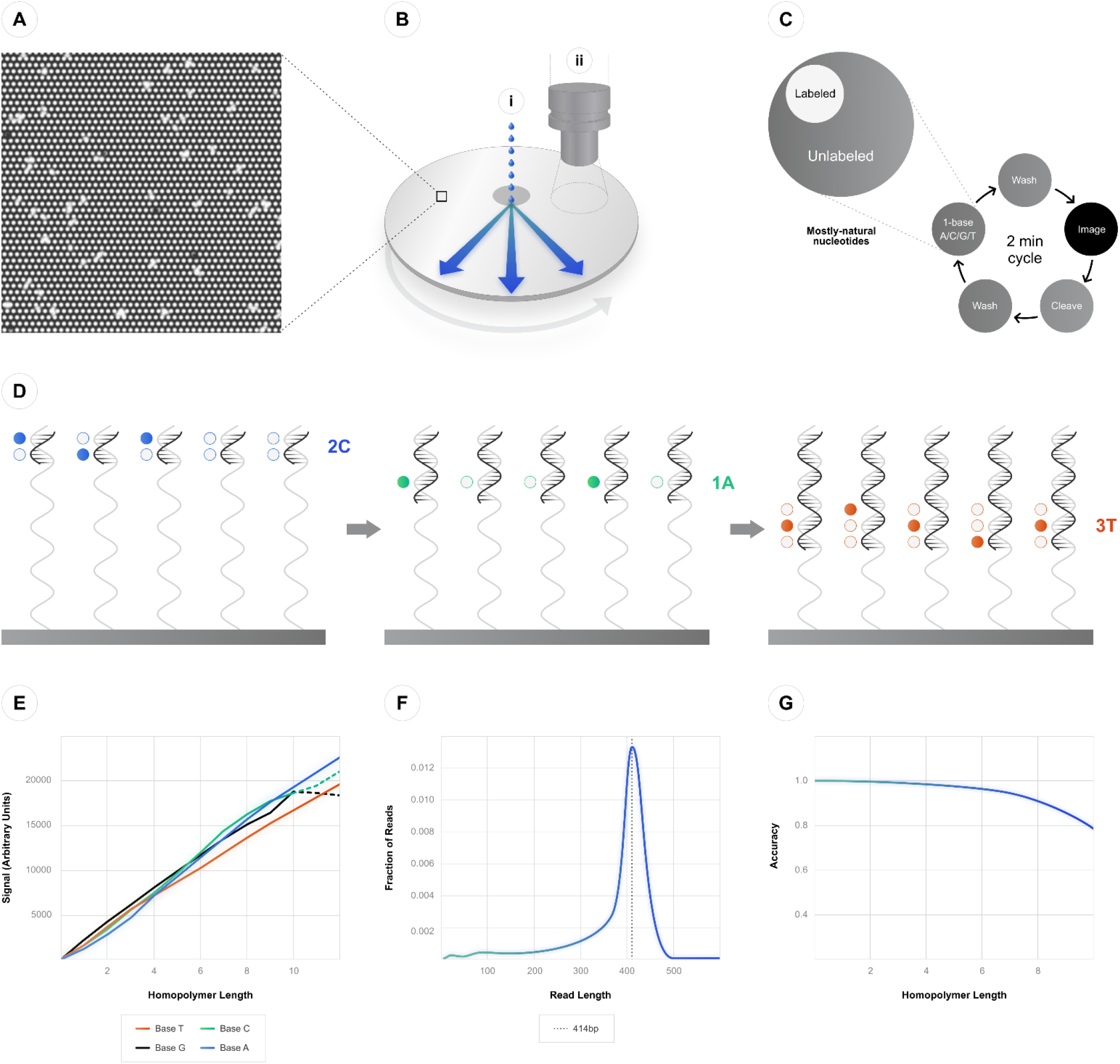
Sequencing Platform. a) Wafer surface patterned at micron resolution to allow binding and sequencing of billions of clonally amplified sequencing beads; b) Open fluidics systems allows (i) dispensing of reagents from dedicated nozzles near the center of the rotating wafer to distribute reagents by centrifugal force and (ii) optical measurement of the entire wafer surface in one continuous step; c) chemistry cycle includes addition of one type of mostly-natural nucleotide mix at a time (dA, dC, dG or dT) followed by imaging and cleavage of the sparse labels; d) In each chemistry cycle, the number of imaged labels is linearly proportional to respective homopolymer length on the template, with minimal number of adjacent fluorophores; e) median signal over 3 different cycles for homopolymer length in the range 0-12, dashed line represent low count of high C/G homopolymers; f) example of prototype run with modal read length >400bp; g) Homopolymer calling accuracy as a function of homopolymer length.

The advantages of the open system are thus several-fold: First, the 200mm wafer is a standard low-cost substrate in the semiconductor industry. Second, the large substrate surface allows deposition of over 10 billion beads at the initial density, with significant headroom to increase density in the future. Third, the spin-dispense system lowers dead-volume and allows efficient reagent delivery, reducing reagent costs. Fourth, the system allows rapid reagent delivery and optical scanning, shortening overall sequencing cycle time. Finally, the “air-gap” between reagent-specific fluidics components leaves minimal opportunity for cross-contamination.

#### Mostly natural sequencing chemistry

The vast majority of sequencing data currently produced for research and clinical use is generated by massively parallel sequencing platforms employing SBS. The most widely used SBS implementation utilizes fluorescently labeled, reversible-terminated nucleotides that are incorporated within a sequencing cycle including all four bases across amplified DNA colonies fixed to a flow-cell. Following completion of this reaction, the entire flowcell is imaged^8^. The main advantages of this approach are high throughput and measurement accuracy, both of which result from the decoupling between the chemistry and optical imaging steps. Another common implementation of SBS introduces one natural (non-terminating) nucleotide base in each cycle and measures the number of incorporated bases in real time using different methods such as pyrosequencing^9^ or changes in pH^10^. The main advantage of the latter approach is that the synthesized DNA is completely unmodified and can therefore be extended with high speed and processivity, even to read lengths of 1kb^11^, while using relatively less costly reagents. However, this approach suffers from higher error rates due to the intrinsic noise of transiently measuring single particles, e.g., protons or photons, emitted during each individual incorporation event.

Mostly natural sequencing-by-synthesis (mnSBS) combines the strengths of these two SBS approaches by utilizing in each sequencing cycle a single base from a mostly natural nucleotide (MNN) mix, which comprises a minority (<20%) of fluorescently labeled, non-terminated nucleotides and a majority of unlabeled, non-terminated (natural) nucleotides (Fig. 1c). The resulting synthesized DNA is mostly unmodified, yet incorporated bases can still be efficiently measured at the reaction endpoint via high throughput optical scanning with a high signal-to-noise ratio, based on hundreds of photons produced by each fluorescent label. At each sequencing cycle clonal beads are exposed to the MNN mix and polymerase extension is performed to incorporate 0, 1, or several bases of a single nucleotide base type (dA, dC, dG or dT) into each growing strand, depending on the length of the respective homopolymer in the corresponding template (Fig. 1d). Following the chemistry step, the entire active surface of the substrate is scanned to measure the fluorescence level of each clonal bead. Then, all fluorescent labels are cleaved and washed, with a typical sequencing cycle length of about 2 minutes. The net benefit of mnSBS is that, in a single sequencing cycle, the total amount of labeled nucleotide incorporated on a clonal bead is linearly proportional to the length of the respective homopolymer in the template, while each individual synthesized DNA strand remains mostly unlabeled – even at longer template homopolymer lengths. mnSBS thereby avoids quenching of fluorescent signals from adjacent labels and instead produces signals proportional to the lengths of homopolymers (Fig. 1e). mnSBS also eliminates most scarring effects of reversible-terminating chemistries, allowing for faster run times and longer read lengths, which we have already demonstrated to surpass 400bp (Fig. 1f). Finally, mnSBS retains a single polymerase on each template strand throughout a sequencing run, significantly enhancing process and cost efficiencies.

#### Neural network-enabled base-calling

Scanning of the entire surface of the wafer generates non-overlapping tile images which are processed using on-board GPUs to extract bead locations and corresponding raw signal vector per clonally amplified sequencing bead over multiple sequencing cycles. Raw signals are proportional to the homopolymer length up to approximately 12 bases. However, raw signals can vary due to multiple factors including dephasing of DNA synthesis in each bead over the sequencing run, spatial differences across the wafer, differing label incorporation rates due to local sequence context, and random sampling noise. To accommodate for systematic variance, we take advantage of recent significant advances in machine learning as well as the large amount of data involved in each run to employ a deep convolutional neural network (CNN) to convert raw signals into sequence reads. The CNN is trained offline on a large, diverse dataset combining data from multiple runs, and it is then recalibrated on a smaller sample of genomic reads from the current run. The run-specific calibration process optimizes for the local variable characteristics of each run, which results in a high level of system robustness and more accurate base calls. The CNN outputs the most likely homopolymer sequence length (i.e., 0-12) for each cycle as well as an estimate of certainty for each base call, which is used to generate base quality scores calibrated for the specific run. Typically, homopolymer calling accuracy is at 99.5% for homopolymer lengths of 1-2 and decreases to 90% at homopolymer lengths of 8 (Fig. 1g). Base substitution error for this type of chemistry is expected to be very low since substitution errors could only be generated as a combination of two or more adjacent homopolymer errors. Following base-calling, data is demultiplexed using inline sample barcodes, and outputted as a standard compressed CRAM file containing the sample information, read sequence, and base quality data, enabling analysis via standard sequence analysis software.

### Sequencing Performance

#### Genome in a Bottle reference samples

To establish sequencing performance, we sequenced the seven standard GIAB reference samples HG001-HG007 ^12^. PCR-free shotgun libraries were generated from the DNA by ligation of DNA adapters including a sample index sequence. Clonally amplified sequencing beads were generated via a fully automated high-scale emulsion PCR system and loaded onto the sequencing wafer. For a full sequencing run, we executed 444 flow cycles to allow reading of the inline sample barcode followed by a single-ended read of length ∼300bp, over a total run time of under 20 hours.

As expected for non-terminating SBS, sequence reads had a distribution of read lengths, with an average of 282bp and mode of 310bp (Fig. 2a). The average amount of sequence generated per sample (after de-duplication) was approximately 60X mean coverage per sample and was down-sampled to 40X for analysis. >99.9% of reads were mappable to Human Genome reference build 38 (GRCh38)^13^ using bwa-mem^14^. The human genome coverage demonstrated good uniformity across the genome with an average F90=1.43 and F95=1.75 (ratio of coverage between the median and the 10^th^ or 5^th^ percentile lowest coverage, respectively) (Fig. 2b). Coverage depth of genomic regions with 20-70% GC content (covering >98% of the human genome) is within a ratio of 0.8-1.2 of the median depth, with lower coverage only at extreme %GC (Fig. 2c). On average, 94.9% and 86.8% of bases had base quality scores of BQ≥20 or BQ≥30, respectively, with high quality maintained throughout the length of the run (Fig. 2d). Measured raw base error rates were <0.1% for high-quality substitution errors (corrected for common SNPs and alignment errors) and 0.3% for total indel errors, with base qualities well-correlated with the observed indel error rate (Fig. 2e, see Methods).

**Figure 2:**
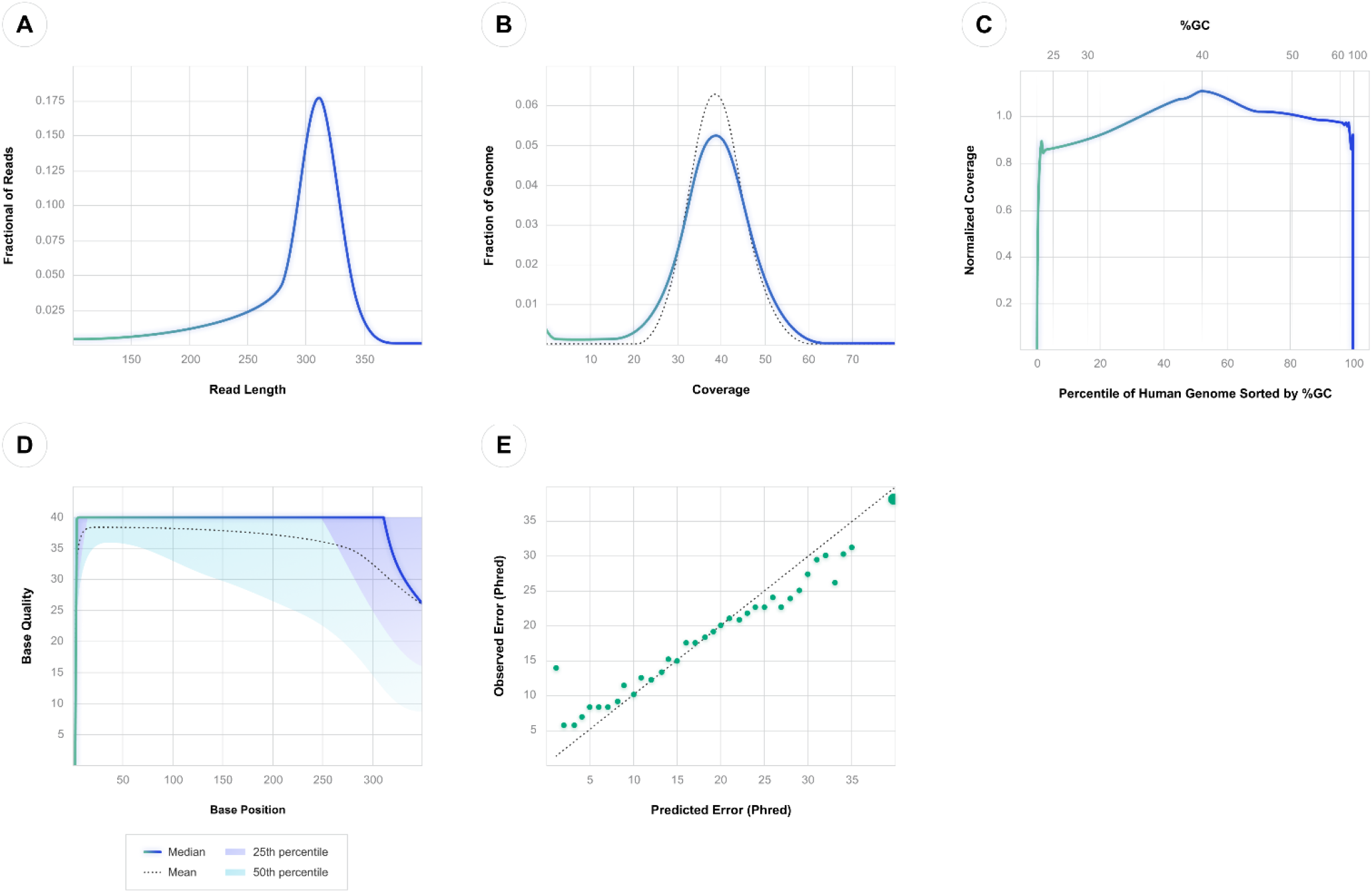
Sequencing Performance for sample HG005. a) Read length distribution; b) Coverage depth distribution across human genome, compared with uniform random sample at same average depth (dashed line); c) Normalized coverage across range of %GC in human genome (HG). Lower x-axis denotes percentile of HG, upper x-axis – the actual %GC value; d) base quality as a function of base position along the read; e) Comparison of predicted and observed homopolymer length classification errors by base quality – see Methods section for details. Larger datapoint contains >85% of the data.

To call germline short variants from WGS data, we modified the GATK HaplotypeCaller algorithm^15^ to account for the unique quality scores that are generated for this data type, by scoring candidate haplotypes using the homopolymer length probabilities represented in the base quality scores. Following variant candidate generation, we scored the variants based on likelihood and filtered them by strand bias and additional variant properties (see Methods).

We assessed the quality of the variant calls by comparing them to reference GIAB truth sets using the respective high-confidence regions^7^ and excluding homopolymer regions of length>10 which are not currently measured by this system (0.4% of the total HCR) (GIAB-HCR). Since low-complexity genomic regions tend to amplify inefficiently, we isolated the sequencing accuracy by further excluding selected low-complexity, tandem-repeats and low mappability regions from the GIAB-HCR while still maintaining 98.2% of the original HCR (UG-HCR, see Methods). The overall concordance of SNP variant calls over the UG-HCR was F1=99.6% (recall=99.7%, precision=99.6%) and of Indel variant calls it was F1=96.4% (recall=96.1%, precision=96.7%). Variant calling accuracy in the GIAB-HCR was F1=99.0% for SNPs and F1=90.6% for Indels, suggesting that a significant fraction of the variant calling errors in this region is indeed related to low-complexity DNA and can likely be improved by optimizing the clonal amplification protocol.

#### Sequencing of 1000 Genomes samples

To demonstrate the scalability of the WGS workflow for large scale studies, we sequenced an additional 224 selected samples from the 1000 Genomes project^16^. We generated an average of 56X mean coverage per sample (after de-duplication) with mean read length of 279bp (mode 302bp) and 86% of bases with BQ≥30. All performance metrics were highly consistent across the full set of samples (Table 1).

**Table 1:** Performance metrics for Genome in a Bottle (GIAB) reference genomes HG001-7, and average performance metrics for 224 additional 1000 Genomes (1000G) reference genomes.

|  | HG001 | HG002 | HG003 | HG004 | HG005 | HG006 | HG007 | GIAB<br>Mean | 1000G<br>Mean | 1000G<br>Std |
| --- | --- | --- | --- | --- | --- | --- | --- | --- | --- | --- |
| **Mean coverage** | 40.05 | 39.58 | 39.67 | 38.56 | 39.16 | 39.53 | 38.32 | 39.27 | 55.65 | 8.14 |
| **% >20X** | 95.9% | 95.7% | 95.9% | 96.6% | 96.8% | 96.1% | 96.7% | 96.2% | 97.8% | 0.4% |
| **% Duplication** | 4.1% | 6.6% | 6.2% | 7.3% | 7.6% | 6.1% | 8.2% | 6.6% | 10.7% | 1.1% |
| **F90\*** | 1.483 | 1.466 | 1.469 | 1.377 | 1.398 | 1.464 | 1.368 | 1.432 | 1.347 | 0.029 |
| **F95\*** | 1.821 | 1.885 | 1.803 | 1.677 | 1.632 | 1.797 | 1.666 | 1.754 | 1.628 | 0.054 |
| **PF reads (M)\*\*** | 469 | 435 | 436 | 433 | 430 | 443 | 433 | 440 | 665 | 103 |
| **% reads aligned** | 99.80% | 99.96% | 99.96% | 99.97% | 99.96% | 99.95% | 99.97% | 99.94% | 99.91% | 0.1% |
| **Mean read length** | 264.8 | 284.9 | 284.4 | 286.7 | 288.6 | 280.3 | 286.4 | 282.3 | 279.5 | 3.5 |
| **Median read length** | 291 | 302 | 302 | 303 | 304 | 301 | 302 | 300.7 | 295.2 | 2.0 |
| **Modal read length** | 309 | 311 | 311 | 311 | 311 | 310 | 311 | 310.6 | 302.3 | 0.6 |
| **% chimeras** | 2.3% | 1.1% | 1.2% | 1.3% | 1.4% | 1.5% | 1.3% | 1.5% | 1.0% | 0.2% |
| **Raw Indel error** | 0.27% | 0.30% | 0.29% | 0.28% | 0.29% | 0.38% | 0.28% | 0.30% | 0.52% | 0.04% |
| **HQ Mismatch error†** | 0.07% | 0.08% | 0.07% | 0.07% | 0.07% | 0.10% | 0.07% | 0.07% | 0.14% | 0.02% |
| **% BQ20 bases** | 95.5% | 94.8% | 95.0% | 95.2% | 95.0% | 93.6% | 95.2% | 94.9% | 94.3% | 0.4% |
| **% BQ30 bases** | 87.3% | 86.4% | 86.8% | 87.6% | 87.1% | 84.6% | 87.7% | 86.8% | 86.5% | 0.8% |
| **Ti/Tv ratio Exome** | 2.97 | 2.89 | 2.95 | 2.90 | 2.98 | 2.93 | 2.97 | 2.94 | 3.06 | 0.03 |
| **Ti/Tv ratio‡** | 2.09 | 2.09 | 2.09 | 2.09 | 2.09 | 2.09 | 2.09 | 2.09 | 2.10 | 0.004 |
| **SNP recall‡** | 99.7% | 99.6% | 99.6% | 99.7% | 99.7% | 99.6% | 99.7% | 99.7% | 99.2% | 0.1% |
| **SNP precision‡** | 99.6% | 99.6% | 99.6% | 99.6% | 99.6% | 99.6% | 99.6% | 99.6% | 99.1% | 0.1% |
| **SNP F1‡** | 99.7% | 99.6% | 99.6% | 99.7% | 99.7% | 99.6% | 99.7% | 99.6% | 99.2% | 0.1% |
| **Indel recall‡** | 96.7% | 96.4% | 96.6% | 95.4% | 96.0% | 95.9% | 96.0% | 96.1% | 95.0% | 0.5% |
| **Indel precision‡** | 97.0% | 96.8% | 97.1% | 96.4% | 97.0% | 96.2% | 96.7% | 96.7% | 94.9% | 0.4% |
| **Indel F1‡** | 96.9% | 96.6% | 96.8% | 95.9% | 96.5% | 96.1% | 96.3% | 96.4% | 95.0% | 0.4% |

\* F90/95: Ratio of coverage between the median and the 10th or 5th percentile lowest coverage, respectively.
\*\* PF: Pass-filter reads. All other metrics were calculated over these reads.
† HQ Mismatch error rate was corrected for germline SNPs and alignment errors (see Methods section).
‡ Variant calling metrics were measured on GIAB HCR excluding long homopolymers and repetitive regions (UG-HCR, see Methods).

For a cohort-level analysis we generated a joint callset from the full set of samples and further scored it based on the combined properties of the cohort^17^ (see Methods). We tested concordance of the variant calls from these samples versus a reference callset^18^ over the intersection of the GIAB HCRs. The average concordance over the entire dataset was F1=99.2% for SNP variant calls and F1=95.0% for Indels in the UG-HCR. Overall Ti/Tv ratio in the genome was measured at 2.09 and 2.94 in the exome.

## Discussion

Most innovation in genomics over the last 20 years is related to the dramatic reductions in the cost of generating large volumes of sequence data. Here, we introduced a novel SBS sequencing architecture, incorporating mostly natural sequencing chemistry and an open fluidics platform, to enable high-throughput sequencing at significantly improved cost efficiency over what is currently available. Beyond the initial implementation, this architecture allows significant headroom for continuous improvements in read length, loading efficiency, bead density, and other factors that will enable significant decrease in cost per Gb over time.

We have demonstrated the accuracy of this system by generating whole-genome data for the seven GIAB reference samples, showing that high sequencing accuracy for SNPs (99.6%) and Indels in homopolymers up to length 10 (96.4%) can be obtained in the vast majority (>98%) of the defined high-confidence regions of these samples. The main accuracy gaps that remain are strongly correlated with repetitive regions and extreme %GC regions that are difficult to amplify efficiently and can likely be minimized by additional optimization of the clonal amplification protocol. Indels in homopolymer regions, which are traditionally the Achilles heel of non-terminating chemistries, can be called with good accuracy up to lengths of 8-10 bases. Further improvements in features such as polymerase fidelity, base calling, and variant calling algorithms, are expected to continue driving overall accuracy gains over time.

We further demonstrated the capacity and stability of our system by sequencing 224 additional reference genome samples. Comparison of the joint variant callset for these samples with reference data demonstrates high similarity in all relevant parameters, with SNP calls being highly concordant (99.2%) with reference data, which is important for allowing the use of new data in combination with legacy datasets.

There are many data-intensive applications which depend upon lower costs to be deployed at scale. For example, studies involving single-cell analysis of millions of cells^19^, high-resolution genomic structure by Hi-C, deep sequencing of cell-free DNA^20^, and whole-genome methylation analysis^21^ have already been demonstrated. Many additional high-throughput sequencing applications, such as deep microbiome analysis, high-resolution spatial profiling, and multi-modal analyses can also be significantly expanded using this platform.

The increase in accessibility and affordability of DNA sequencing over the last two decades has already profoundly changed the way biological and clinical research are carried out. It is exciting to imagine how this new level of low-cost sequencing will enable continued advances, and how the expanded availability of large-scale genomic data will facilitate routine clinical sequencing and implementation of precision medicine across all areas of human health.

## Methods

### Sample preparation and Sequencing

The Broad Institute Genomics Platform created PCR-free Ultima Genomics (UG) libraries in an automated 96-well format using the Agilent Bravo automated liquid handling platform. Starting from 250 ng of genomic DNA, samples were prepared using the NEBNext® Ultra™ II FS DNA Library Prep Kit. Genomic DNA was enzymatically fragmented for 10 minutes. After end-repair and A-tailing, custom UG-specific adapters containing molecular barcodes were ligated. This was followed by a double-sided SPRI-based size selection cleanup to target fragments averaging 550-600bp. In-process quality control checks were carried out in the form of an automated electrophoresis (Agilent Bioanalyzer) to ensure proper fragment size, and quantitative-PCR (qPCR) to measure the final library concentration. Using a Hamilton Starlet automated liquid handler, samples were pooled volumetrically according to qPCR results such that each sample received equal representation in the final pool. Sample pools were then seeded onto UG sequencing beads, pre-enriched, and amplified by emulsion PCR, leveraging UG’s automated sequencing bead preparation system (AMP1). The material generated by the UG AMP1 system was subjected to QC before loading onto the sequencer. Sequencing was performed on two UG100 sequencing systems, using early access version of UG sequencing chemistry. The flow-based sequencers were run for 444 flow-cycles (111 cycles across each of the four nucleotides [T, G, C, A]).

### Base calling

Base calling was performed using a CNN that outputs the most likely homopolymer sequence length (i.e., 0-12) for each cycle as well as an estimate of certainty for each base call. The output of the base-calling network per read is a matrix of size (# of flows, 13) where position (*h, f*) in the matrix describes the probability that the true homopolymer length corresponding to the read’s flow *f* is *h*. The read sequence in the output file is generated as the most likely homopolymer length call in each flow, and the probabilities of the alternate homopolymer lengths (errors) are encoded in the quality string and the optional tp and t0 tags. For additional information, see CRAM format in Supplementary Materials.

GIAB samples were base-called with base-calling pipeline version APL4.6, quality filtered by read quality (rq≤0.7) and down-sampled to 40X. 1000 Genomes samples were base-called with base-calling pipeline version APL4.2.

To calculate the predicted vs observed error rates (Fig. 2e), 1% of the reads were randomly sampled and encoded base qualities were converted back to flow space. Errors in homopolymer length classification vs reference were tallied and compared to the quality estimates.

### Performance Metrics

Performance metrics listed in Table 1 were calculated using GATK Picard^22^ tools: AlignmentSummaryMetrics, RawWgsMetrics, and QualityYieldMetrics.

To most accurately estimate substitution error rate based on available WGS 1% of the reads were randomly sampled and filtered to chromosome 9. Only reads with high mapping quality (≥60) were selected, and positions corresponding to known population variants in dbSNP were ignored (for GIAB samples, the positions corresponding to the ground truth were ignored). We also removed errors that are likely due to local misalignment of indels (the more common error mode) by ignoring positions that are close to the read ends (<=5bp) or with an adjacent mismatch.

### Variant Calling

Variant calling was performed using a version of GATK HaplotypeCaller^15^ algorithm that was updated to use the flow-space error estimations from the base-calling network to calculate the likelihood of having sequenced a given read from a candidate haplotype, in place of the native PairHMM algorithm that assumes mismatches as the predominant error mode. In addition, variants in homopolymers of length 12 or above were ignored, and an iterative candidate pruning scheme was implemented to generate the most accurate variant calls. For additional information, see GATK release details in Supplementary Materials.

Variant call filtering was implemented using a random forest classifier retrained per sample using an approximate labeling scheme. 200,000 variants were selected and labeled as likely positive, if they matched dbSNP, and likely false if they matched a database of common artifacts generated over 200 unrelated samples. The input features of the classifier were the parameters of the variant from the GATK output (e.g. quality, strand bias, local motif etc.). The trained classifier was applied to all variant calls to produce the final filtered VCF.

For joint calling on the 231 sample dataset, individual variant calls (GVCF) were integrated into GenomicsDB as part of the GATK^17^ with additional novel annotations included for filtering. The single sample scores were averaged across sites and used as an annotation along with the average number of assembled haplotypes and filtered haplotypes within HaplotypeCaller, as well as Read Position Rank Sum, Fisher Strand, Strand Odds Ratio, and Qual by Depth in an Isolation Forest model that filtered the sites at the Joint Callset level. Two filtering thresholds were chosen (one for SNPs and one for Indels) by using one of the GIAB samples that was included in the callset to determine a max F1 score. This threshold was used to filter the entire callset.

### Variant Calling Performance Evaluation

Variant calling performance (recall, precision F1), was calculated using vcfeval^23^ to compare single sample variant callset (vcf) with GIAB truth set (v4.2.1)^7^ for reference samples HG001-7 or WGS reference callset^18^ for the remaining samples.

The evaluation region was defined as the corresponding GIAB high-confidence region (HCR v4.2.1), with the following exclusions [% of full HCR]:

*GIAB-HCR* (total 99.6% of full HCR):

- Homopolymer regions of length 11 and higher + 4 flanking bases [0.4%]

*UG-HCR* (total 98.2% of full HCR):

- Homopolymer regions of length 11 and higher + 4 flanking bases [0.4%]
- AT-rich regions: all 40 bp regions with 95% or higher AT content [0.3%]
- Short tandem repeats regions, with specific defined thresholds [0.3%]
- Low mappability and coverage: 50bp regions with mean mappable coverage>15X in 90 independent samples [1.1%]

BED files containing specific exclusion areas are available as supplementary material.

## Acknowledgements

Ultima Genomics acknowledges early support from NHGRI via grants R44HG010558, R44HG011060.

## Supplementary materials

Supplementary data is available at: https://www.ultimagenomics.com/publications/

